# The topology of interpersonal neural network in weak social ties

**DOI:** 10.1101/2023.04.09.536147

**Authors:** Yuto Kurihara, Toru Takahashi, Rieko Osu

## Abstract

People often have opportunities to engage in social interactions with strangers, which have been reported to contribute to their well-being. Although strategies for social interaction between strangers differ from those between acquaintances, the differences in neural basis of social interaction have not been fully elucidated. In this study, we examined the geometrical properties of interpersonal neural networks in pairs of strangers and acquaintances during joint tapping using dual electroencephalography (EEG). Twenty-one pairs of participants performed antiphase joint tapping under four different conditions. Intra-brain synchronizations were calculated using the weighted phase lag index (wPLI) for all possible intra-brain pairs of the 29 channels (_29_*C*_2_ = 406), and inter-brain synchronizations were calculated using the phase locking value (PLV) for all possible inter-brain pairs of the 29 channels (29 × 29 = 841) in the theta, alpha, and beta frequency bands. Electrode pairs with larger wPLI and PLV than their surrogates were defined as the nodes (EEG channels) and edges (connections between nodes) of the neural networks. We then calculated the global efficiency, local efficiency, clustering coefficient, and modularity derived from graph theory for the combined intra- and inter-brain networks of each pair. No significant differences in the tapping phase variance were identified between the stranger and acquaintance pairs. However, in the combined intra- and inter-brain theta EEG (4–7 Hz) networks, stranger pairs showed larger local efficiency and cluster coefficients than acquaintance pairs, indicating that the two brains of stranger pairs were more densely connected. Moreover, in the beta EEG bands, the modularity of the two brains was low in the fast condition, indicating that the two brains were coupled when the task demand was high. Our results show that weak social ties promote more extensive social interactions and result in dense brain-to-brain coupling.

## 1. Introduction

In everyday life, we experience different levels of intimacy in interpersonal relationships with acquaintances, friends, romantic partners, or strangers. Previous studies have suggested that interacting with people with whom one has stronger social ties, such as romantic partners, family members, and close friends, maybe beneficial (Kiecolt-Glaser & Wilson, 2017; Meyer & Sledge, 2020; Robles et al., 2014; Slatcher & Schoebi, 2017). People tend to experience less loneliness after intimate interactions (Wheeler et al., 1983). Since the intensity and quality of friendships are positively correlated with life satisfaction (Amati et al., 2018), strong social relationships are expected to contribute to well-being. However, in our daily lives, we have many opportunities for social interaction outside our close social groups, such as with strangers or acquaintances with whom we have weak social ties. Interestingly, interacting with people with weak social ties has also been found to contribute to well-being (Dunn et al., 2007; Granovetter, 1973; Sandstrom & Dunn, 2014). Sandstrom and Dunn (2014) demonstrated that simply engaging in social interactions with a coffee shop barista (a stranger) can increase people’s sense of belonging and well-being.

Although many studies have examined neural responses during intimate interactions, almost no studies have examined interactions in weak social ties. In general, pairs with intimate interpersonal relationships, such as romantic couples and parents, tend to have more synchronized brain activities than pairs with non-special interpersonal relationships. For example, romantic couples (male-female pairs) have been found to have a higher correlation between brain-to-brain electroencephalography (EEG) spectra than stranger pairs (male-female pairs) in face-to-face conversations (Djalovski et al., 2021; Kinreich et al., 2017). Romantic partners also have higher inter-brain synchronization (functional near-infrared spectroscopy, fNIRS) than friend pairs during two-handed touching (Goldstein et al., 2018; Long et al., 2021), and cooperation tasks (Djalovski et al., 2021; Duan et al., 2022; Pan et al., 2017). Parent-child pairs have been found to have higher neural synchrony (fNIRS) than stranger-adult and child-pair pairs during cooperation tasks (Kruppa et al., 2021; Reindl et al., 2018, 2022). These studies indicate that intimate pairs exhibit high interbrain synchronization. In contrast, in the empathy giving task (shareing their events each other), Djalovski et al. (2021) revealed that weak-tie pairs (strangers and friends) showed higher inter-brain EEG synchronization with lower behavioral synchronization than romantic couples, while they showed lower inter-brain EEG synchronization with lower behavioral synchronization than romantic couples in their motor task. Our study also showed that weak-tie pairs (including stranger and acquaintance pairs) showed a negative correlation between behavioral synchrony and inter-brain EEG synchronization (Kurihara et al., 2022). From these studies, it can be postulated that the relationship between intimacy and inter-brain synchronization is inversely correlated (i.e. complementary) for weak-social tie pairs (Djalovski et al., 2021). Although social interactions between same-gender individuals are experienced on a daily basis, almost no study to date has compared neural synchrony between same-gender strangers and acquaintance pairs. If we extrapolate the results of intimate relationships, such as romantic or parent-child pairs, to those of weak-tie relationships, we may hypothesize that neural synchrony is higher in acquaintance pairs than in stranger pairs owing to their degree of intimacy. However, if intimacy and neural synchrony are complementary, we hypothesized that neural synchrony would be higher in stranger pairs than in acquaintance pairs. Therefore, in this study, we examined which hypothesis was supported.

Hence, we compared the interpersonal neural networks between stranger and acquaintance pairs during anti-phase joint tapping tasks. Although the task selected is a motor task, it requires the prediction of the movement of the partner. Hence, this task was selected. In addition, compared with in-phase tapping, the anti-phase mode is advantageous in avoiding spurious inter-brain EEG synchronization caused by similarities in movement across individuals. We conducted tests with four tapping conditions: slow (requested inter-tap interval [ITI]:0.5 s), fast (requested ITI:0.25 s), free-speed (preferred ITI), and pseudo (no interaction). To quantify neural synchrony, we performed 29-channel electroencephalography (EEG) and applied a graph-theoretical approach to characterize the topology of the multi-brain network, in addition to comparison using the synchronization indices [Phase Locking Values (PLV) / weighted Phase Lag Index (wPLI)] among EEG channels. For each pair, we created a binary undirected graph consisting of nodes (EEG channels) and edges (connections between nodes). The topology of the interpersonal neural network can detect the state changes within pairs, such as the demand for musical coordination in guitar duets (Sänger et al., 2012), social coordination in romantic couples (Müller & Lindenberger, 2014), the degree of flight cooperation in pilot pairs (Toppi et al., 2016), the difference between joint and solo actions (Astolfi et al., 2020), and emotional communication in parent-child pairs (Santamaria et al., 2020). However, to the best of our knowledge, whether the neural network topology differs when the relations of the pairs differ has not been verified. Thus, we aimed to verify whether neural network topology is related to interpersonal relationships using graph theory. To reveal the properties of the network topology for weak-tie pairs, we recruited stranger pairs (meeting each other for the first time in an experiment) and acquaintance pairs (one participant brought his/her partner). To evaluate the synchronization strength, we calculated the wPLI (Vinck et al., 2011) for intra-brain electrode pairs and PLV (Lachaux et al., 1999) for inter-brain electrode pairs. To test the topology, we compared the edge number, global efficiency, path length, local efficiency, clustering coefficient, and modularity as graph-theoretical measures between stranger/acquaintance groups and tapping conditions.

## 2. Methods

### 2.1. Participants

A total of 27 pairs of right-handed participants (13 male pairs, 14 female pairs; mean age=22.38 years, SD=2.94 years) were enrolled in the study. Two male and four female pairs were excluded from the analysis due to recording artifacts. Thus, 21 pairs were included in the analysis. Of these 21 pairs, 10 already knew each other before participating in the experiment (defined as acquaintance pairs), while the remaining 11 pairs met for the first time in this experiment (defined as stranger pairs). The experimental procedures were approved by the Ethical Review Committee of Waseda University and conducted in accordance with the code of ethics of the world medical association (Declaration of Helsinki) for experiments involving humans. All the participants provided written informed consent. The data used in this study were obtained from our previous study (Kurihara et al., 2022). Specifically, data from 13 pairs (strangers: five pairs, acquaintances: eight pairs) were obtained from Kurihara et al. (2022).

### 2.2. Behavioral Task

Participants in each pair were seated side by side (Fig.1a) and asked to gaze at a fixation point during the tapping task (Fig.2b). The distance between the participants was approximately 70 cm. Participants were asked to coordinate anti-phase tapping using two computer mice. Each computer mouse was placed on two tables (40 × 50 cm). The participant who tapped first used the mouse’s left click with his/her right index finger. The tapping produced sound feedback (sound frequency of 440 Hz). Each participant listened to the sounds produced by their own taps and their partner’s taps through earphones.

**Fig.1.**
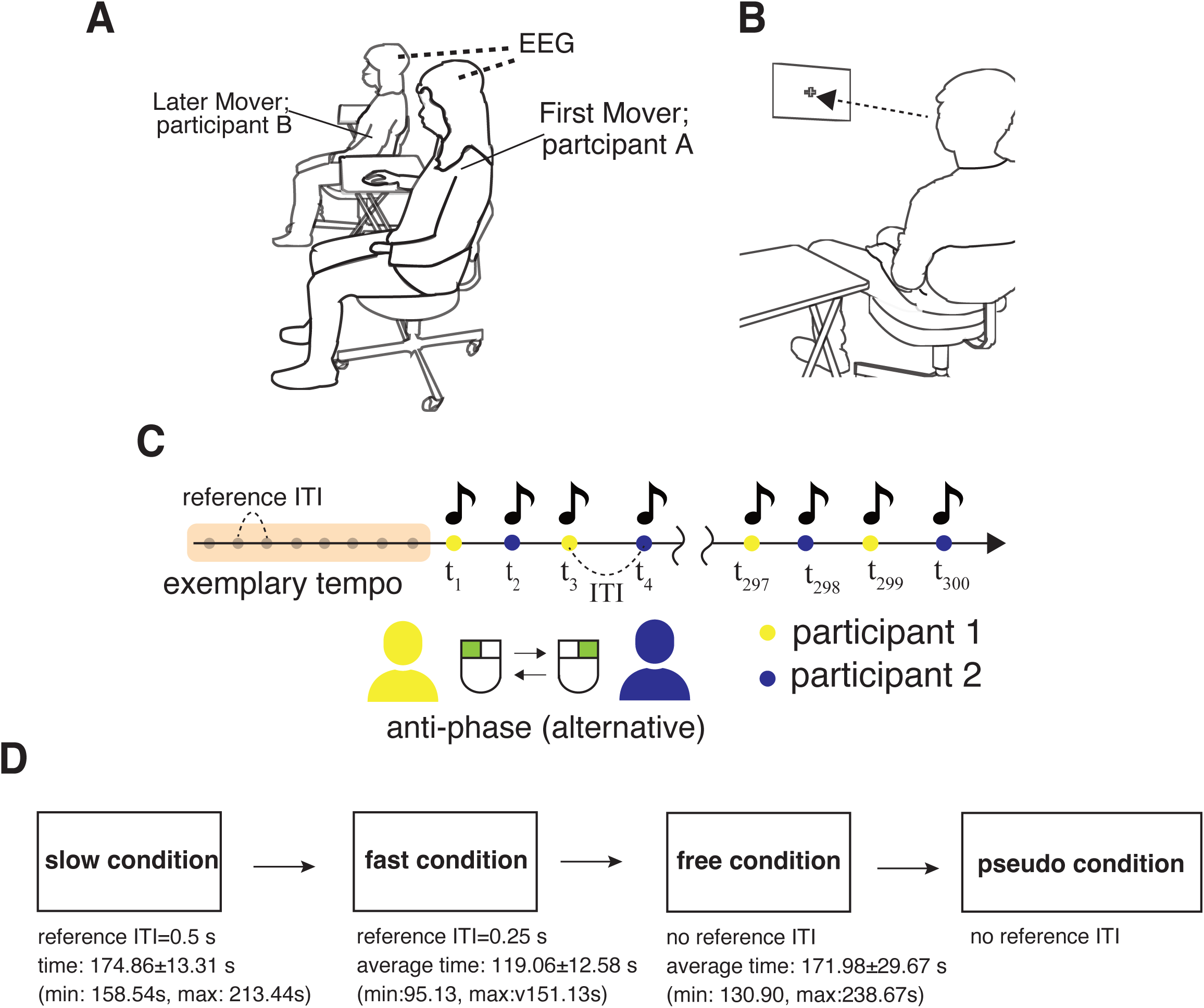
Experimental setups. (a) Participants sat side-by-side and wore two wireless electroencephalographs (EEG). (b) Each participant gazed at a fixation point in front of him/her during anti-phase tapping. (c) The participants performed alternative (anti-phase) tapping. (d) We conducted four different tapping conditions: slow, fast, free, and pseudo conditions.

The participants participated in four sessions of anti-phase tapping, with each participant performing 150 taps per session (300 taps per pair). The tapping sessions consisted of four conditions: slow, fast, free, and pseudo conditions. In the slow and fast conditions, the participants listened to an exemplary frequency from a metronome, as a reference tempo. After the first eight sounds were transmitted, the metronome was switched off and the pairs started tapping at a frequency as close as possible to the memorized reference Inter-tap interval (ITI) (slow:0.5s, fast:0.25s). In the free condition, there was no reference for ITI and the participants tapped at their preferred frequency. In the pseudo condition, after eight sounds of ITI=0.50s, the participants continued tapping the metronome (ITI = 0.50s).

### 2.3. EEG acquisition

We acquired EEG activities from each participant simultaneously. The EEG device had a 29-channel acquisition system (Quick-30; Cognionics, San Diego, CA, USA) in accordance with the international 10/20 system: Fp1/Fp2, AF3/AF4, F3/F4, F7/F8, FC5/FC6, C3/C4, T7/T8, CP5/CP6, P3/P4, P7/P8, PO3/PO4, PO7/PO8, O1/O2, Fz, Cz, and Pz (Fig.2a). The sampling rate was 500 Hz. A reference electrode was placed on the left earlobe.

**Fig.2.**
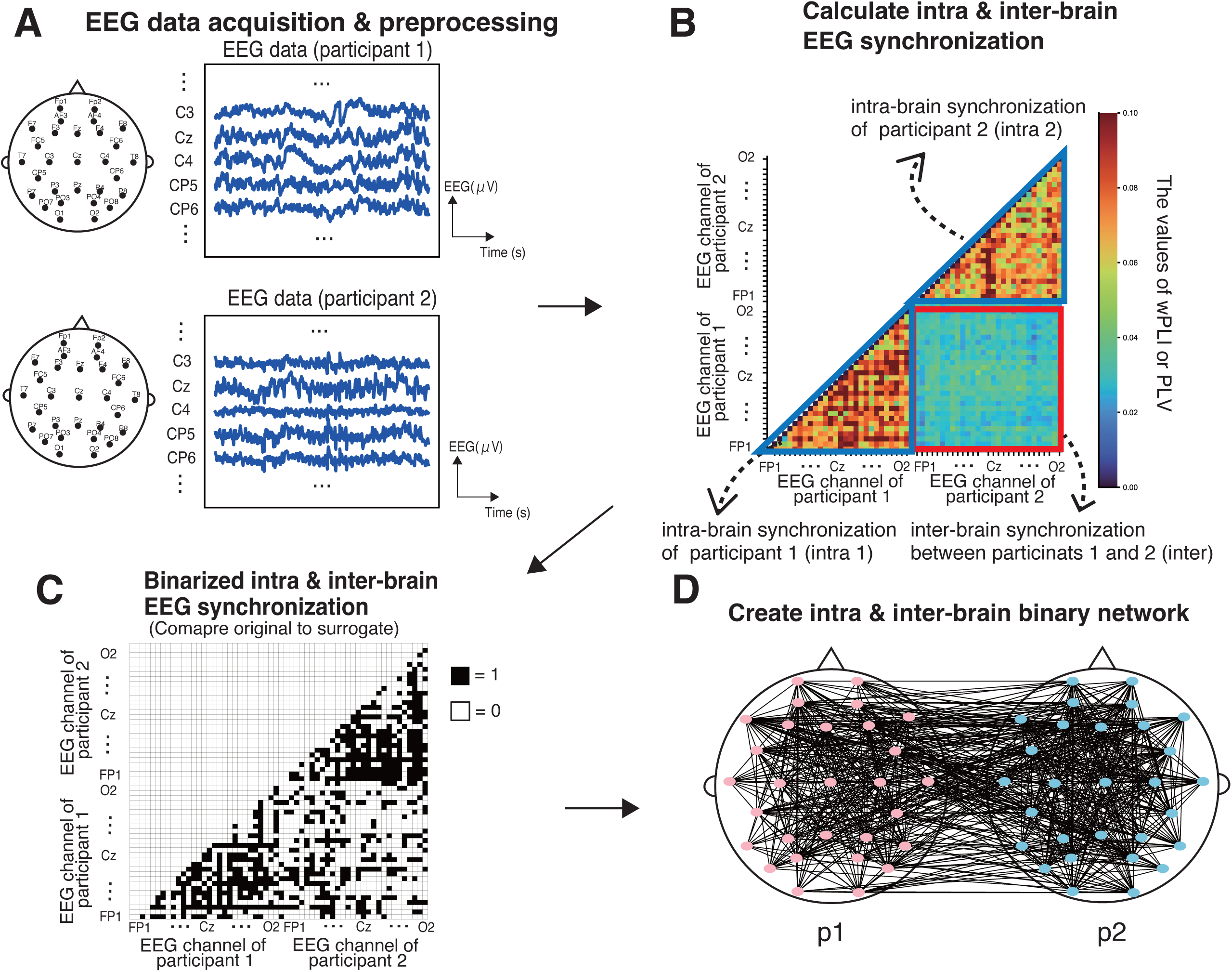
The procedure of intra- and inter-brain electroencephalography (EEG) analysis. (a) We collected EEG data from two participants simultaneously. These EEG datasets were filtered into theta, alpha, and beta bands. (b) We calculated intra- and inter-brain EEG synchronization. The area enclosed by the yellow square shows the intrabrain synchronization of participants 1 and 2. The “intra 1” shows within brain synchronization in participant 1 and the “intra 2” shows within brain synchronization of participant 2. The area enclosed by the red square shows inter-brain synchronization. (c) We compared original EEG synchronization to surrogate EEG synchronization and conducted binarization of intra- and inter-brain synchronization (values equal 1 if original synchronization was significantly larger than that of surrogate, else, it equals 0). (d) From the binary matrices of intra- and inter-brain, we created an interpersonal neural network.

### 2.4. Behavioral analysis

We computed the variance from the anti-phase to confirm whether the tapping phase variance differed between the stranger and acquaintance pairs. First, we calculated the relative phase (RP) of tapping using a circular measure to confirm whether the tapping coordination was antiphase (RP=180°). (Kodama et al., 2015; Kurihara et al., 2022; Novembre et al., 2017). The RP was defined using tapping times:

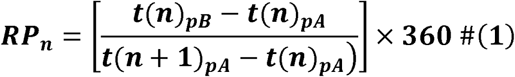

where *t*(*n*)*_pA_* and *t*(*n*)*_pB_* denote the tapping time (second) of participants A and B, respectively. Here, n denotes the number of tapping. The RP ranges from 0°to 360°.

We calculated the circular standard deviation of RP (SDRP) as follows:

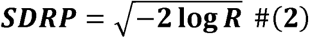

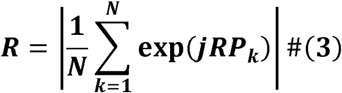

where SDRP represents the instability of anti-phase tapping. The R is the resultant length of the average RP vector. The N is the total number of tapping data points. The j represents the complex number.

### 2.5. EEG preprocessing

The EEG data were downsampled to 250 Hz and then filtered to eliminate artifacts using a bandpass filter with a range of 1–45 Hz. Visual checks were performed to eliminate problematic EEG channels. The analysis was omitted for bad channels (mean number of bad channels for all participants: 2.42; SD: 1.25). We performed an independent component analysis (ICA) of the EEG to decrease or remove artifacts (electrooculogram, muscular noise, perspiration, and movement). To extract the theta (4–7 Hz), alpha (8–12 Hz), and beta (13–30 Hz) frequency bands, EEG data were band-pass filtered. EEG preprocessing was conducted using MNE Python (0.20.7).

### 2.6. Synchrony of each EEG channel pair in the intra-brain

The wPLI was used to estimate the intrabrain synchrony of each EEG channel pair (Fig.2b). wPLI measures the extent to which the difference in the phase angle between two signals is distributed towards the positive or negative parts of the imaginary axis in the complex plane (Imperatori et al., 2019; Vinck et al., 2011). The wPLI enables the estimation of intra-brain synchronization without the deleterious impact of volume conduction and avoids pseudo-synchronization (Cohen, 2015; Yokoyama et al., 2018). The wPLI ranges from 0 (unsynchronized) to 1 (perfectly synchronized). The wPLI equation is as follows:

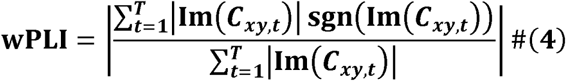

where *C_xy,t_* is the cross-spectrum density between signals x(t) and y(t) at time t. The T is the total number of sampling points. x(t) and y(t) show the EEG oscillations filtered into the theta, alpha, or beta frequency bands. The Im{ · } shows the imaginary operator. Phase angles x(t) and y(t) were determined using the Hilbert Transform. We calculated the wPLI in each EEG channel across the other 28 channels (i.e., the total number of channel pairs was _29_*C*_2_ = 406) for each tapping condition (slow, fast, free, and pseudo-condition) and EEG frequency (theta, alpha, and beta bands).

### 2.7. Synchrony of each EEG channel pair in the inter-brain

We used PLV to estimate the interbrain synchrony of each EEG channel pair (Czeszumski et al., 2020; Lachaux et al., 1999) (Fig.2b). The PLV measures the phase synchrony between the signals. The PLV was used for each pair (i,k) of electrodes between participants A and B. The PLV_i,k_ (synchronization between channel i and k) equation is as follows:

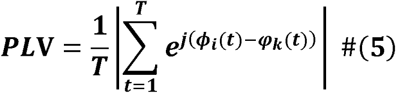

where *φ_i_*(*t*) and *φ_k_*(*t*) are the phase angles of the EEG channels of Participants A and B, respectively. T is the total number of EEG sample points. j denotes a complex number. The phase angles were acquired using the Hilbert Transform. The PLV ranged from 0 (unsynchronized) to 1 (perfectly synchronized). We calculated the inter-brain synchronization in each EEG channel of the participants across 29 channels (i.e., the total number of channel pairs was 29 × 29 = 841).

### 2.8. Creation of surrogate datasets and construction of binary undirected graphs from intra- and inter-brain networks

To create binary undirected graphs with channel pairs that have synchrony, the wPLI or PLI of each channel pair was compared with the surrogate and those with values significantly above the chance level were extracted (Santamaria et al., 2020; Toppi et al., 2016). To generate surrogate data, we applied a Fourier transform to each EEG data for each channel and performed a random permutation of phase values in the frequency domain while maintaining the power of each frequency and applied an inverse Fourier transform (Santamaria et al., 2020). For each individual, EEG channel, and tapping condition, 100 surrogate datasets were created to produce a distribution of the wPLI or PLV values for significance testing. The wPLI or PLV values obtained from the original data were then compared with those from the surrogate data using Welch’s t-test (all p-values were adjusted using the Bonferroni correction). EEG channel pairs with significantly larger wPLI or PLV values than their surrogates were selected as EEG channels for an undirected graph and connected by the edges of value 1 (Fig.2c).

### 2.9. Graph Theory

Binary undirected graphs were created for each participant pair and each tapping condition, utilizing the selected nodes (EEG channels) and edges (binary connections between nodes) from the comparison between the original and surrogate. To synthetically describe the features of functional networks synthetically, we employed multiple measures of functional brain network topology (Stone et al., 2019). We calculated graph-theoretical indices for the combined intra- and inter-brain networks (Fig.3e). Specifically, we calculated the edge number, Global Efficiency, Path Length, Local Efficiency, Clustering Coefficient, and Modularity. These graph-theoretical indices were calculated using Network X (version 3.4) (Hagberg et al., 2008).

*Edge number* was defined as the number of edges in the intra- and inter-brain graphs. The equation of the Edge number *EN* is as follows:

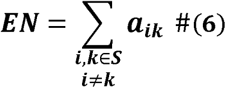

where *a_ik_* indicates the connection status between nodes *i* and *k*: *a_ik_* = 1 when nodes *i* and *k* are connected and *a_ik_* = 0 when they are not connected. S represents the set of all nodes (EEG channels) in the intra- and inter-brain networks.

*Global Efficiency* is the average inverse shortest path length between all the pairs of nodes in a network (Astolfi et al., 2020; Latora & Marchiori, 2001; Rubinov & Sporns, 2010). The equation for the global efficiency GE is as follows:

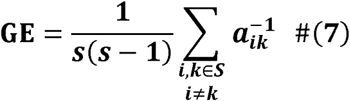

where the *s* is the total number of nodes (Rubinov & Sporns, 2010).

Path length (PL) is the network’s average shortest path length (Rubinov & Sporns, 2010; Watts & Strogatz, 1998). The shortest path length between two nodes is the shortest number of edges that must be traversed from one node to the next (Sporns et al., 2004). The equation for the path length PL is as follows:

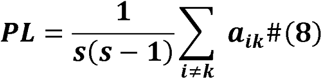

Local Efficiency represents the efficiency of communication between all the nodes around node *i* in a network (Astolfi et al., 2020; Latora & Marchiori, 2001; Rubinov & Sporns, 2010). The equation of local efficiency *LE* is as follows:.

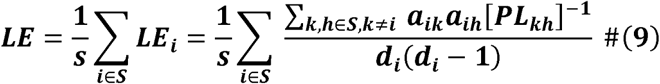

where *LE_i_* is the local efficiency of node *i*. *PL_kh_* is the length of the shortest path between nodes *k* and *h,* which contains only the neighbors of node *i*. *d_i_* is the number of links connected to a node.

*The clustering coefficient (CC)* expresses the degree of connectedness between a node’s neighbors. It is defined as the number of triangles that surround a node or the number of neighbors that are neighbors of each other (Astolfi et al., 2020; Rubinov & Sporns, 2010; Watts & Strogatz, 1998). The equation for the CC is as follows:

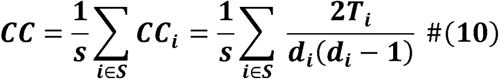

where *CC_i_* is the clustering coefficient of node *i*. *T_i_* is the number of triangles of the three nodes around node *i* and *d_i_* is the number of edges in node *i*.

*Modularity i*s the degree to which a network can be separated into a module of nodes with a maximally possible number of within-module links and a minimally possible number of between-module links (Arenas et al., 2007; Newman, 2004, 2006; Rubinov & Sporns, 2010). We defined each of the two participant networks as a subnetwork community. Thus, higher modularity indicates that the two participant-identified subnetworks do not communicate with each other in the brain. Modularity equation Q is as follows:

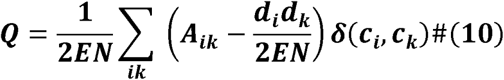

where EN is the number of edges, *A* is the adjacency matrix of the graph, and *k_i_* is the degree of node *i*. *d_i_* shows the number of edges connected to a node *i*. *δ*(*c_i_*, *c_j_*) is 1 if node *i* and *k* are in the same community else it is 0.

### 2.10 Statical Analyses

We performed a two-way analysis of variance (ANOVA) on the average of intra-brain synchronization (averaged-wPLI), the average of inter-brain synchronization (averaged-PLV), Edge number, Global Efficiency, Path Length, Local Efficiency, Clustering Coefficient, and Modularity for the effect of partner (stranger and acquaintance pairs) and tapping conditions (slow, fast, free, and pseudo conditions).

### 2.11 Data availability statement

The EEG and tapping data generated or analyzed during this study are included in the published article.

## 3. Results

### 3.1. Behavioral analysis results

The means and standard deviations of the tapping variance (SDRP) for each interpersonal relationship and tapping conditions are reported in Table 1. We conducted Welch’s t-test for SDRP between stranger and acquaintance pairs in slow, fast, and free conditions. There were no significant differences in SDRP between stranger and acquaintance pairs in the slow-, fast-, or free-tapping conditions (slow: *t*_17.45_ = -0.084, *p_adj_* = 1.0, *d* = -0.037; fast: *t*_13.91_ = -0.637, *p_adj_* = 1.0, *d* = -0.285; free: *t*_13.91_ = -0.717, *p_adj_* = 1.0, *d* = -0.315; all p-values were adjusted by Bonferroni correction).

**Table 1:** The mean and standard deviation of SDRP.

|  | slow |  | fast |  | free |  |
| --- | --- | --- | --- | --- | --- | --- |
|  | Mean | SD | Mean | SD | Mean | SD |
| stranger | 17.175 | 8.152 | 29.501 | 21.033 | 17.175 | 8.152 |
| acquaintance | 14.459 | 9.121 | 38.028 | 37.315 | 14.459 | 9.121 |

SD, standard deviation.

### 3.2. The average brain synchronization strength

#### 3.2.1 Intra-brain synchronization

First, we calculated the average wPLI (intra-brain synchronization) for each participant (defined as *averaged-wPLI*) in the theta, alpha, and beta frequency bands. The overall number of objects analyzed was 42 (21 pairs × 2). Average intra-brain synchronization was calculated using the synchronization strengths of all channels. Next, we conducted a two-way ANOVA on averaged-wPLIs with strangers/acquaintances as a between-group effect and within the four tapping conditions for the average of intra-brain synchronization in the theta, alpha, and beta frequency bands.

For the theta frequency bands, we found a significant main effect of stranger/acquaintance groups (*F*_1,40_= 4.9257, *p* = 0.0322, 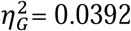, *Mean_stranger_*=0.0752, *SD_stranger_*=0.021, *Mean_acquaintance_* =0.0657, *SD_acquaintance_*=0.026). Thus, stranger pairs showed a higher averaged-wPLI than acquaintance pairs in this frequency band. There were no significant main effects of tapping conditions (*F*_3,120_= 1.4087, *p*= 0.2436, 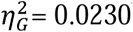) or interaction effect (*F*_3,120_= 0.3925, *p* = 0.7586, 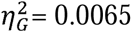). In the alpha frequency bands, there were no significant main effects of groups (*F*_1,40_= 0.5145, *p* = 0.4774, 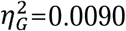), the main effect of tapping condition (*F*_3,120_= 0.8771, *p* = 0.4551, 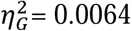) or their interaction effect (*F*_3,120_= 0.2741, *p* = 0.8440, 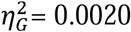). In the beta, there were no significant main effects of groups (*F*_1,40_= 2.7512, *p* = 0.1050, 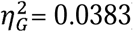), the main effect of the tapping condition (*F*_3,120_= 0.7848, *p* = 0.5047, 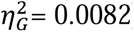), or their interaction effect (*F*_3,120_= 0.8016, *p*= 0.4954, 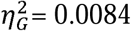). Thus, stranger pairs have greater intra-brain synchronization than acquaintance pairs in the theta bands.

#### 3.2.2 Inter-brain synchronization

First, we calculated the average inter-brain synchronization (PLV) for each participant (defined as *averaged-PLV*) in the theta, alpha, and beta frequency bands. A total of 21 participant pairs were analyzed. We also conducted a two-way ANOVA of the averaged PLV between stranger/acquaintance groups and within the four tapping conditions for the average inter-brain synchronization in the theta, alpha, and beta frequency bands. In the theta frequency band, we confirmed marginally significant main effect of stranger/acquaintance groups (*F*_1,19_= 3.6857, *p*= 0.0700, 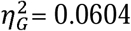), significant main effect of tapping conditions (*F*_3,57_*=* 7.80 54, *p* = 0.0002, 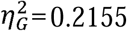), and no significant interaction effect (*F*_3,57_= 1.4369, *p* = 0.2415, 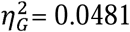). Post hoc paired t-tests using Holm correction showed that the average inter-brain synchronization in the fast condition was higher than that in the slow, free, and pseudo conditions (slow vs. fast: *t*_19_ = 2.6557, *p_adj_* = 0.0468, *d* = 0.823; fast vs. free: *t*_19_= 3.8895, *p_adj_* = 0.0059, *d* = 1.142; fast vs. pseudo: *t*_19_ = 3.6922, *p_adj_* = 0.0059, *d* = 0.909). Post-hoc paired t-tests of other combinations of tapping conditions showed no significant differences (slow vs. free: *t*_19_ = 1.4185, *p_adj_* = 0.5167, *d* = 0.418; slow vs. pseudo: *t*_19_= 0.3057, *p_adj_* = 0.7631, *d* = 0.083; free vs. pseudo: *t*_19_= 1.2861, *p_adj_* = 0.5167, *d* = -0.357).

In the alpha frequency band, the factor of tapping conditions showed a significant main effect (*F*_3,57_= 11.1775, *p*<. 001, 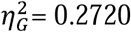). The factor of stranger/acquaintance groups showed no main effect (*F*_1,19_= 0.2031, *p* = 0.6573, 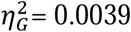). In addition, there was no interaction effect (*F*_3,57_= 0.3169, *p* = 0.8131, 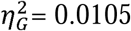). Post-hoc paired t-tests using Holm correction showed that the averaged-PLV of the fast condition was higher than that in the slow, free, and pseudo conditions (slow vs. fast: *t*_19_= 5.1466, *p_adj_* =0.0003, *d* = 1.288; fast vs. free: *t*_19_= 4.1226, *p_adj_* = 0.0017, *d* = 1.211; fast vs. pseudo: *t*_19_ = 3.6522, *p_adj_* = 0.0051, *d* = 1.223). Post-hoc paired t-tests of other combinations of tapping conditions showed no significant differences (slow vs. free: *t*_19_ = 0.1622, *p_adj_* = 1.00, *d* = 0.039; slow vs. pseudo: *t*_19_ = 0.5498, *p_adj_* = 1.00, *d* = 0.198; slow vs. free: *t*_19_= 0.4619, *p_adj_* = 1.00, *d* = 0.143).

For the beta bands, the tapping condition showed a significant main effect (*F*_3,57_= 17.2520, *p*<. 001, 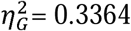). The factor of stranger/acquaintance groups showed no main effect (*F*_1,19_= 0.5105, *p* = 0.4836, 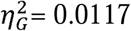). In addition, there was no interaction effect (*F*_3,57_= 1.5 3 5 9, *p*= 0.2150, 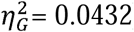). Post hoc paired t-tests using Holm correction showed that the average inter-brain synchronization of the fast condition was higher than that of the slow, free, and pseudo conditions (slow vs. fast: *t*_19_ = 5.9184, *p_adj_* = 0.0001, *d* = 1.221; fast vs. free: *t*_19_= 5.1736, *p_adj_* = 0.0002, *d* = 1.419; fast vs. pseudo: *t*_19_= 4.2965, *p_adj_* = 0.0012, *d* = 1.328). Post hoc paired t-tests of other combinations of tapping conditions showed no significant differences (slow vs. free: *t*_19_ = 0.2447, *p_adj_* = 1.00, *d* = 0.076; slow vs. pseudo: *t*_19_= 0.3050, *p_adj_* = 1.00, *d* = 0.083; slow vs. free: *t*_19_ = 0.9189, *p_adj_* = 1.00, *d* = 0.227). Thus, these results showed that the fast condition of average inter-brain synchronization was the highest among the four tapping conditions in the theta, alpha, and beta frequency bands. In addition, stranger pairs had slightly higher average inter-brain theta band synchronization than acquaintance pairs did.

### 3.3. The geometric features of neural networks

The averages of the total number of edges in the combined intra- and inter-brain networks for strainger pairs and acquaintance pairs were reported in Table 3 for each tapping condition and EEG frequency band. We conducted a two-way repeated ANOVA (2 × 2 factorial design: interpersonal relationship and tapping condition) for edge number, global efficiency, path length, local efficiency, clustering coefficient, and modularity of combined intra- and inter-brain networks in the theta, alpha, and beta frequency bands. The results showed a significant main effect of partner for local efficiency (*F*_1,19_= 4.630, *p* = 0.0445, 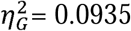) and *clustering coefficient* (*F*_3,57_= 5.091, *p* = 0.036, 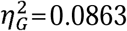; Fig.5a, b) in the theta frequency band. As shown in Figures 5a and 5b, stranger pairs showed higher local efficiencies and clustering coefficients than acquaintance pairs did. There were no significant main effects of tapping conditions for local efficiency (*F*_3,57_= 0.3942, *p* = 0.7577, 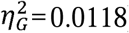) or clustering coefficient (*F*_3,57_= 0.0992, *p* = 0.960, 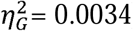), and no significant interaction of partner and tapping conditions for local efficiency (*F*_3,57_= 1.616, *p* = 0.196, 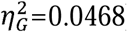) or clustering coefficient (*F*_3,57_= 1.043, *p* = 0.381, 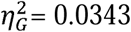) in the theta frequency band. In addition, we confirmed the significant main effect of tapping conditions for modularity (*F*_3,57_= 3.638, *p* = 0.018, 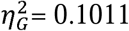) in the beta frequency band. The post hoc analysis of modularity for the tapping condition showed that the fast condition was marginally significantly higher than the slow and free conditions (slow vs. fast: *t*_19_= 2.7710, *p_adj_* = 0.0730, *d* = 0.827; fast vs. free: *t*_19_ = 2.3921, *p_adj_* = 0.0817, *d* = 0.665; fast vs. pseudo: *t*_19_= 1.4939, *p_adj_* = 0.0817, *d* = 0.481). Post hoc paired t-tests of other combinations of tapping conditions showed no significant differences (slow vs. free: *t*_19_ = 0.8695, *p_adj_* = 0.7909, *d* = 0.196; slow vs. pseudo: *t*_19_= 1.4967, *p_adj_* = 0.4527, *d* = 0.351; free vs. pseudo: *t*_19_ = 0.7675, *p_adj_* = 0.7909, *d* = 0.169). The other graph theory indices (edge number, global efficiency, and path length) showed no significant differences in the theta, alpha, and beta frequency bands, respectively (Table A1). Thus, stranger pairs indicated higher local efficiency and clustering coefficients than acquaintance pairs at theta frequencies. We also confirmed the lowest modularity under fast-tapping conditions.

**Fig. 5.**
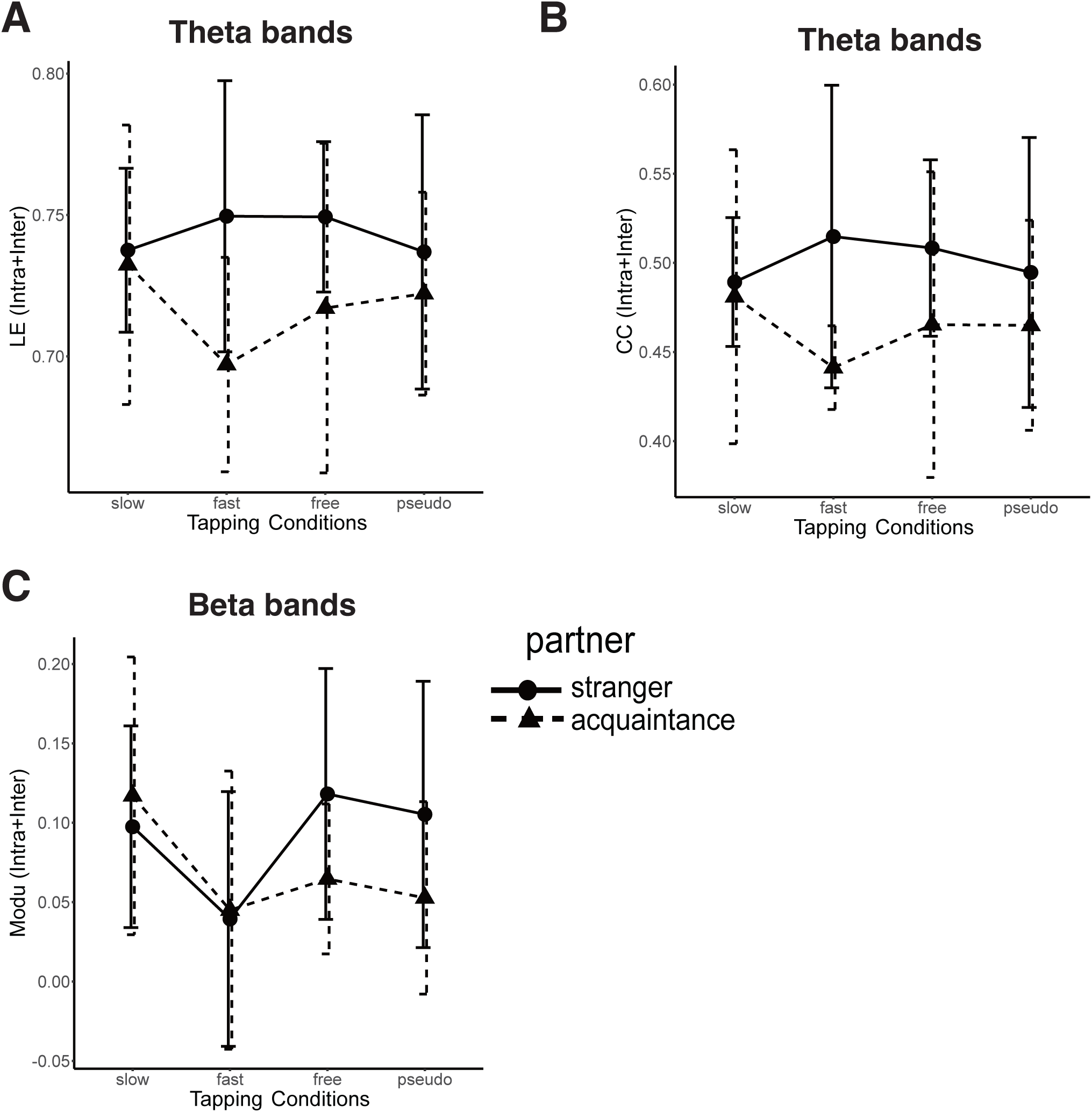
The line plots of graph theory indices of intra- and inter-brain networks in the partner and tapping conditions. (a) There was a significant main effect of interpersonal relationship for Local Efficiency (LE) of intra- and inter-brain theta EEG network. (b) There was a significant main effect of interpersonal relationship for Clustering Coefficient (CC) of intra- and inter-brain theta EEG network. (c) There was a significant main effect of tapping conditions for Modularity (Modu) of intra- and inter-brain beta EEG network. These error bar means standard deviation.

**Table 3:**
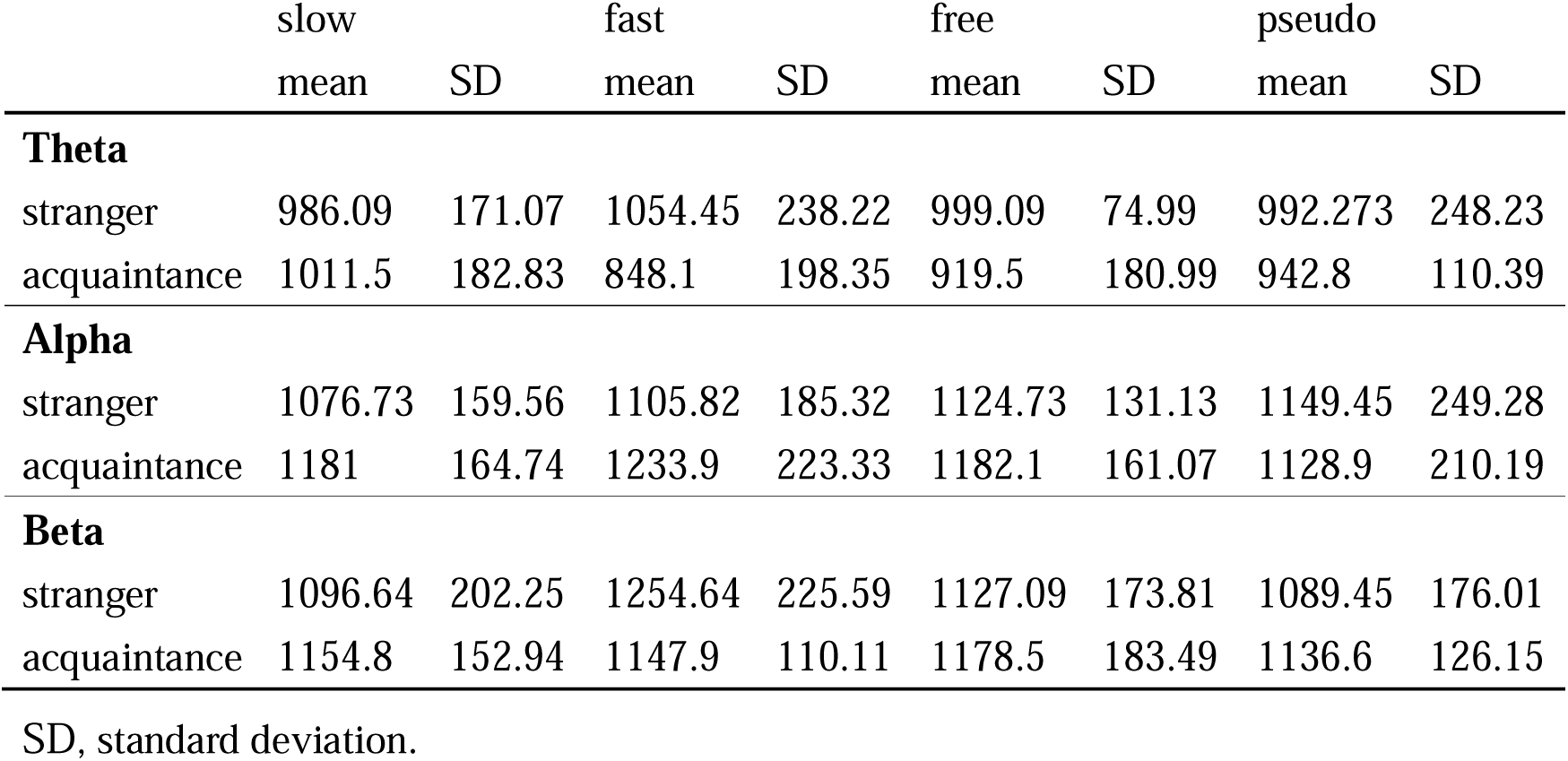
The mean and SD of the edge number in the combined intra- and inter-brain network.

|  | slow<br>mean | SD | fast<br>mean | SD | free<br>mean | SD | pseudo<br>mean | SD |
| --- | --- | --- | --- | --- | --- | --- | --- | --- |
| **Theta** |  |  |  |  |  |  |  |  |
| stranger | 986.09 | 171.07 | 1054.45 | 238.22 | 999.09 | 74.99 | 992.273 | 248.23 |
| acquaintance | 1011.5 | 182.83 | 848.1 | 198.35 | 919.5 | 180.99 | 942.8 | 110.39 |
| **Alpha** |  |  |  |  |  |  |  |  |
| stranger | 1076.73 | 159.56 | 1105.82 | 185.32 | 1124.73 | 131.13 | 1149.45 | 249.28 |
| acquaintance | 1181 | 164.74 | 1233.9 | 223.33 | 1182.1 | 161.07 | 1128.9 | 210.19 |
| **Beta** |  |  |  |  |  |  |  |  |
| stranger | 1096.64 | 202.25 | 1254.64 | 225.59 | 1127.09 | 173.81 | 1089.45 | 176.01 |
| acquaintance | 1154.8 | 152.94 | 1147.9 | 110.11 | 1178.5 | 183.49 | 1136.6 | 126.15 |

SD, standard deviation.

## Discussion

In this study, we examined the differences in intra- and inter-brain synchronization and interpersonal neural network topology between stranger and acquaintance pairs during a joint-tapping task. There was no difference in the behavioral performance indicated by the variance of the tapping phase between the two groups. However, we found significant differences in the average strength of intra-brain synchronization and network topology between the two groups. Specifically, the neural networks of stranger pairs were more densely connected, as indicated by higher local efficiency and clustering coefficient values. These findings suggest that stranger pairs may engage in a higher level of social interaction than acquaintance pairs during tasks requiring mutual prediction.

We found that the stranger pairs were more densely connected to each other in the neural network than acquaintance pairs. Previous research has suggested that pairs with strong social ties, such as romantic partners and parent-child pairs, show greater brain synchronization than pairs with weaker social ties (e.g., strangers) (Djalovski et al., 2021; Kinreich et al., 2017; Long et al., 2021; Pan et al., 2017; Reindl et al., 2018). These results suggest that pairs with higher levels of intimacy are more considerate of others and engage in higher levels of social interaction, resulting in greater brain synchronization. On the other hand, other lines of studies, have shown that the efficiency and integration of brain-to-brain networks depend on task properties, and if the task requires higher social interaction between participants, the two brains are more synchronized (Astolfi et al., 2015, 2020; Falk & Bassett, 2017). These results indicate that brain-to-brain synchronization depends on the social demands of the task, rather than on the intimacy level of the interacting individuals. Therefore, it is conceivable that some tasks require stronger interactions between strangers, resulting in higher synchronization than those between acquaintances. In fact, Djalovski et al. (2021) showed that in social tasks, inter-brain synchronization is higher in stranger pairs than in friends or couples (Djalovski et al., 2021). Our findings suggest that strangers with relatively weak social ties may engage in higher levels of social interaction than acquaintances with medium or strong social ties. There may be a nonlinear relationship between the level of intimacy and social engagement, and thus, inter-brain synchronization.

The reasons for the brains of stranger pairs being more synchronized with each other than those of acquaintance pairs when interaction demand is high could be explained based on previous studies. An fNIRS hyperscanning study has suggested that social interactions with a high mental load induce higher inter-brain synchrony (Park et al., 2022). Furthermore, in socially attentive situations, more mentalization is required, which may induce brain synchronization between each other (Clark & Dumas, 2015; Nozawa et al., 2019). A previous study has shown that cooperation between strangers may produce high mental loading (P. Sariñana-González et al., 2017; Patricia Sariñana-González et al., 2019). Thus, it is reasonable that stranger pairs were more densely connected to each other in the neural network than acquaintance pairs since stranger pairs were more attentive to the mutual prediction of their behavior than acquaintance pairs. Kikuchi et al. argued that cooperative stranger pairs may be more synchronized in their brains than acquaintance pairs who are less cooperative in economic exchange tasks than stranger pairs (Kikuchi et al., 2022).

We found that interpersonal relationships were associated with lower frequencies (the theta band). It has been shown that the more complex the social interaction, the greater the EEG activity in the theta band (Blume et al., 2015). Blume et al. interpreted brain activity in the theta band as reflecting increased demands on the participants’ attention and working memory resources when observing complex social interactions. In fact, many studies have shown that theta EEG oscillations play a role in cognitive functions, such as memory encoding and retrieval, working memory retention, novelty detection, and realizing the need for top-down control (Cavanagh & Frank, 2014; Cavanagh et al., 2010, 2012; Itthipuripat et al., 2013; Jacobs et al., 2006; Rutishauser et al., 2010). For stranger pairs, the partner is a more novel social stimulus than for acquaintance pairs. A load of top-down processing and retention of working memory may be higher in the process of tapping when the partner is a stranger than when the partner is an acquaintance (P. Sariñana-González et al., 2017; Patricia Sariñana-González et al., 2019). Thus, it is reasonable to find an increase in theta-band synchrony related to working memory and top-down processing in stranger pairs.

We found that the beta frequency modularity of the two brains in the fast condition was lower than that in the other tapping conditions. The lower modularity indicated that the two brains were coupled rather than independent when higher interpersonal coordination was required, which is consistent with our previous study showing larger EEG synchronization during fast tapping (Kurihara et al., 2022). This could be attributed to the high difficulty of the joint-tapping task in the fast condition, which induced the coupling of the two brains. As mentioned above, fNIRS hyperscanning studies have shown that the degree of inter-brain synchronization is higher when the cooperative tapping task is more difficult (Park et al., 2022). The target tempo of joint tapping in the fast condition was more difficult to stably coordinate than in the slow condition (standard ITI=0.50s), making it a more challenging task. To achieve stable interpersonal coordination at a fast tempo, it is necessary to predict each other’s movements more accurately. This mutual prediction could be an important factor in brain-to-brain synchronization (Hamilton, 2021).

Previous studies have indicated that beta oscillations are modulated by rhythmic motor control (Bourguignon et al., 2019; Neuper & Pfurtscheller, 2001; Salenius et al., 1997; Salmelin & Hari, 1994). Additionally, auditory-motor rhythm learning is related to beta neural circuits (Edagawa & Kawasaki, 2017). Rosso et al., (2022) revealed that EEG beta power plays a role in interpersonal motor synchronization (finger-tapping tasks). They suggested that EEG beta power may be a critical mechanism underlying interpersonal synchronization, possibly enabling mutual predictions between coupled individuals and leading to the co-regulation of timing and overt mutual adaptation. Using transcranial alternating current stimulation (tACS), (Novembre et al., 2017) elicited beta-band oscillations over the left motor cortex in pairs of participants who performed a joint finger-tapping task. They revealed the induced phase coupling of beta-band neural oscillations in the motor cortex of two individuals, facilitating the establishment of synchronized movement through interpersonal alignment of sensorimotor processes controlling rhythmic movement initiation. Thus, it is reasonable to assume that EEG oscillations are modulated by joint tapping.

Our study has two limitations. Firstly, the sample analyzed was limited to 21 dyads (stranger pairs:11 pairs, acquaintance pairs:10 pairs). Notwithstanding the sample dimension, we detected a difference in the network topology between stranger and acquaintance pairs. To confirm these results further, a larger sample size is required. Secondly, in graph theory analysis, there were only a limited number of methods for selecting important EEG channel pairs. Our selection of significant EEG channel pairs (in comparison with surrogate datasets) to create a neural network for each pair was based on a previous study (Santamaria et al., 2020). There are possible ways of channel selection in group-level analysis, such as selecting EEG channel pairs whose synchronization indices are significantly higher in target tasks (e.g., joint tapping) than in baseline tasks (e.g., independent tapping or resting); however, these do not apply to generating a network for each participant since it requires a statistical test for each channel pair of each participant (Kawasaki et al., 2018; Kurihara et al., 2022).

## Conclusion

To the best of our knowledge, this is the first study comparing the geometric structures of neural networks across interpersonal relationships and coordination. We conducted a two-factor experiment in which the interpersonal levels involved stranger and acquaintance and the levels of interpersonal coordination were slow, fast, free, and pseudo-antiphase joint tapping conditions. We found that, in the EEG theta band, the two brains of stranger pairs were more connected than in acquaintance pairs. In addition, in the beta EEG bands, the networks were more connected in the fast condition, where the difficulty of interpersonal coordination was higher than in the other conditions. Our results show that weak social ties require more extensive social interactions, resulting in dense brain-to-brain coupling.

## Credit Author Statement

**Yuto Kurihara:** Conceptualization, Data curation, Formal Analysis, Investigation, Methodology, Software, Visualization, Writing – original draft, writing – review, and editing. **Toru Takahashi**: Conceptualization, Methodology, Writing, review, and editing. **Rieko Osu:** Conceptualization, Funding acquisition, Methodology, Supervision, Validation, Writing, review, and editing

## Acknowledgements

We would like to thank S. Okazaki for his experimental programming of tapping task. And We would like to thank everyone who took part in the study.

