## Supplementary material for "The topology of interpersonal neural network in weak social ties": Table A1

|  | Interpersonal relationship | |  | Tapping Conditions | |  | Interaction |  |  |
| --- | --- | --- | --- | --- | --- | --- | --- | --- | --- |
|  | F (1,19) | p | Effect size | F (3,57) | p | Effect size | F (3,57) | p | Effect size |
| **Theta** |  |  |  |  |  |  |  |  |  |
| EN | 2.961 | 0.102 | 0.048 | 0.462 | 0.710 | 0.016 | 1.127 | 0.346 | 0.039 |
| GE | 2.715 | 0.116 | 0.044 | 0.492 | 0.689 | 0.017 | 1.206 | 0.316 | 0.041 |
| LE | **4.630** | **0.045** | **0.093** | 0.394 | 0.758 | 0.011 | 1.616 | 0.196 | 0.047 |
| CC | **5.091** | **0.036** | **0.086** | 0.099 | 0.960 | 0.003 | 1.043 | 0.381 | 0.034 |
| Trans | 3.601 | 0.073 | 0.064 | 0.320 | 0.811 | 0.011 | 1.215 | 0.313 | 0.039 |
| PL | 2.306 | 0.145 | 0.039 | 0.544 | 0.654 | 0.019 | 1.349 | 0.268 | 0.045 |
| Modu | 0.113 | 0.741 | 0.002 | 0.564 | 0.641 | 0.020 | 1.134 | 0.343 | 0.039 |
| **Alpha** |  |  |  |  |  |  |  |  |  |
| EN | 0.993 | 0.332 | 0.027 | 0.172 | 0.915 | 0.004 | 0.549 | 0.651 | 0.014 |
| GE | 1.016 | 0.326 | 0.027 | 0.179 | 0.910 | 0.005 | 0.527 | 0.665 | 0.013 |
| LE | 0.229 | 0.638 | 0.008 | 0.197 | 0.898 | 0.004 | 0.237 | 0.871 | 0.005 |
| CC | 0.113 | 0.741 | 0.004 | 0.257 | 0.856 | 0.004 | 0.305 | 0.822 | 0.005 |
| Trans | 0.055 | 0.818 | 0.002 | 0.476 | 0.700 | 0.009 | 0.603 | 0.616 | 0.011 |
| PL | 1.059 | 0.317 | 0.027 | 0.203 | 0.894 | 0.005 | 0.486 | 0.693 | 0.013 |
| Modu | 0.037 | 0.851 | 0.001 | 1.000 | 0.400 | 0.026 | 0.317 | 0.813 | 0.008 |
| **Beta** |  |  |  |  |  |  |  |  |  |
| EN | 0.008 | 0.932 | 0.0002 | 0.574 | 0.635 | 0.017 | 1.249 | 0.301 | 0.035 |
| GE | 0.006 | 0.938 | 0.0001 | 0.575 | 0.634 | 0.017 | 1.234 | 0.306 | 0.035 |
| LE | 0.120 | 0.733 | 0.003 | 0.356 | 0.785 | 0.009 | 0.853 | 0.471 | 0.021 |
| CC | 0.264 | 0.613 | 0.007 | 0.458 | 0.713 | 0.011 | 1.060 | 0.373 | 0.025 |
| Trans | 0.449 | 0.511 | 0.012 | 0.281 | 0.839 | 0.007 | 1.435 | 0.242 | 0.034 |
| PL | 0.004 | 0.950 | 0.0001 | 0.578 | 0.632 | 0.017 | 1.206 | 0.316 | 0.35 |
| Modu | 0.925 | 0.348 | 0.020 | **3.638** | **0.018** | **0.101** | 1.737 | 0.170 | 0.051 |

**Table A1:** The results of two-way repeated ANOVA (2 × 2 factorial design: interpersonal relationship and tapping condition) for edge number (EN), global efficiency (GE), local efficiency (LE), clustering coefficient (CC), path length (PL), and modularity of combined intra- and inter-brain networks in the theta, alpha, and beta frequency bands.

Note Effect size: generalized eta squared
